# Stoichiometric Aquatic Food-Web Models Coupling Pelagic and Benthic Zones

**DOI:** 10.64898/2026.09.07.749907

**Authors:** Ramiro Ramirez

## Abstract

Ecological stoichiometry allows us to investigate how food quality affects food-web population dynamics. We consider periphyton, phytoplankton, and a shared *Daphnia* consumer in a closed system with separate benthic and pelagic phosphate pools. The producers have variable phosphorus-to-carbon ratios, while the consumer maintains a constant ratio. We construct the model, establish boundedness under stated conditions, characterize its grazer-free equilibrium families and their local spectra, derive conditions for positive consumer growth, and examine its numerical bifurcation structure as light-dependent carrying capacity varies. The bifurcation diagrams contain stable and unstable equilibrium branches together with a stable periodic family, revealing a more complex response to light enrichment than a single sequence of steady states and oscillations. A descending consumer equilibrium branch illustrates that greater light availability need not increase consumer biomass. We then introduce periodically varying carrying capacities and compare weak and strong seasonal forcing. Stronger forcing produces brief consumer rebounds in the high-light example, although the peaks diminish over the displayed interval. Together, the growth conditions and branch diagrams provide a framework for interpreting population dynamics in terms of food quantity and food quality. Independent numerical continuation identifies a Hopf crossing and a fold; the termination mechanism of the periodic branch remains unresolved.

## 1 Introduction

Light and nutrients are fundamental components that affect the primary production and quality of an aquatic system. In shallow lakes, extensive littoral zones provide growth potential for periphyton. Periphyton contributes biomass and affects nutrient uptake and transfer between organisms in both the benthic and pelagic zones [1, 2]. Its quality as food can therefore affect the growth of consumers and the linkage between these habitats. Many factors influence periphyton quality, including light, nutrients, temperature, and grazing pressure [1, 3].

Phytoplankton provides another food source near the surface of the water. High phytoplankton densities can decrease light penetration, affecting periphyton at the bottom of the aquatic system. Food-web structure and nutrient enrichment can thus influence both producers and the connection between benthic and pelagic processes [1, 4]. A consumer that feeds on both producers also connects the two habitats through its diet and nutrient recycling.

The light:nutrient hypothesis explains how the balance of light and phosphorus can affect producer food quality [5]. High supplies of light relative to phosphorus tend to produce phosphorus-poor biomass, whereas lower light relative to phosphorus can produce more phosphorus-rich biomass. Consequently, an increase in producer biomass does not necessarily lead to an increase in consumer growth. Producer biomass can be limited by light or nutrients, and the phosphorus-to-carbon ratio of that biomass determines its quality as food. These relationships have been investigated in stoichiometric producer–grazer models [6, 7, 8].

Explicitly tracking producer phosphorus and free nutrients provides a way to study how uptake and storage affect these relationships [9]. Earlier work also considered enrichment in a stoichiometric system with two producers and one consumer [10], and light-dependent seasonality in a producer–grazer system [11]. Recent research has developed these ideas through habitat coupling, food selection, and environmental variation.

For habitat coupling, the placement of organisms and nutrients affects which resources are available to each producer. Borrelli and Relyea [12] reviewed spatial structure in freshwater food webs, emphasizing the need to represent connections within lakes. Muniz-Júnior et al. [13] modeled phytoplankton, periphyton, and zooplankton to investigate habitat coupling and stability in shallow lakes. Sinha et al. [14] examined a coupled benthic–pelagic ecosystem with four living compartments and two nutrient compartments, while Zhang et al. [15] studied spatially asymmetric competition for light and nutrients. More directly related to the present producer subsystem, Yan et al. [16] considered variable producer quotas and separate habitat nutrient pools. Their formulation does not include a dynamic grazer population. These studies motivate representing nutrient location and producer competition together with explicit grazer food-quality feedback.

A shared grazer introduces a second connection between the habitats because its growth depends on the foods it consumes. Ahmed et al. [17] studied stoichiometric foraging in a phytoplankton–periphyton–*Daphnia* system. Their analyzed reduction uses rapidly adjusting nutrient uptake to express producer quotas algebraically. McConnell’s 2026 dissertation [18] considers two producers, a shared consumer, and two dynamic quotas, with one common free nutrient pool and prey-specific feeding responses. These are close comparisons for the present model, which retains separate benthic and pelagic pools and limits consumer production by the quality of the mixed diet. Adaptive feeding has also been studied in a single producer–grazer system by Oladepo and Peace [19] and in two consumers sharing one resource by Oladepo and Peace [20]. Those behavioral extensions provide context for the fixed feeding parameters used here.

Environmental variation affects these interactions through both food quantity and food quality. Building on earlier seasonal formulations, Sun et al. [21] studied seasonal light in a stoichiometric phytoplankton–zooplankton model, including boundary and persistence dynamics. Related work treats periodic nutrient–algal thresholds [22] and changing carrying capacity with environmental noise [23]; the latter concerns environmental change rather than regular seasonal forcing. In a marine setting, Matsumoto et al. [24] examined food-quality-dependent grazing on two phytoplankton groups under seasonal changes in ocean structure. These studies make seasonal forcing and multiple food sources established parts of the background to the present habitat-coupled model.

Recent work also shows why the timing of food quality deserves attention. Twining et al. [25] discussed nutritional mismatches under climate change, and Anderson et al. [26] showed how nutrient storage links past thermal exposure to current phytoplankton growth. Temperature–stoichiometry interactions provide a related perspective: Sentis et al. [27], first published online in 2021, examined biomass distribution and stability, while Diehl et al. [28] combined experiments and modeling to investigate a warming-induced shift associated with poor food quality. Subsequent studies examined temperature-dependent nutritional demands [29] and the interaction of light, nutrients, and phytoplankton thermal responses [30, 31]. These findings motivate attention to nutrient adjustment and environmental timing, but they do not establish a storage-memory or thermal mechanism in the light-only model considered here.

In this study, we consider periphyton, phytoplankton, and *Daphnia* in a system with separate benthic and pelagic phosphate pools. We assume that phytoplankton can take up phosphate from both pools, while periphyton takes up phosphate from the benthic pool. We first construct the constant-light model and then add light-dependent seasonality. Our main objective is to examine how light enrichment changes the equilibrium and periodic branches of this coupled food web, and how seasonal forcing changes the associated population trajectories. The formulation makes the habitat-specific uptake and recycling pathways explicit. Determining whether those pathways change consumer dynamics relative to the closely related models above requires a controlled comparison and remains outside the present analysis. We present the model construction in Section 2, establish boundedness under the stated conditions in Section 3, analyze equilibria and local stability in Section 4, derive consumer growth and invasion conditions in Section 5, examine the bifurcation diagrams and population trajectories in Section 6, and discuss their biological interpretation in Section 7.

## 2 Model construction

### 2.1 State variables and assumptions

Suppose we have a fixed amount of phosphate, *P*_*tot*_, in a closed aquatic system with a consumer, *y*, and two autotrophs: periphyton, *w*, and phytoplankton, *x*. Let *Q*_*x*_ and *Q*_*w*_ denote the producers’ phosphorus-to-carbon (P:C) ratios, and let *θ*_*y*_ denote the consumer’s constant P:C ratio. We write the dissolved phosphate concentrations in the benthic and pelagic zones as *P*_*B*_ and *P*_*p*_, respectively. All concentrations use a common model volume; separate habitat volumes and transport between them are not explicitly resolved.

We use the following assumptions.

1. The total mass of phosphorus in the entire system is fixed; the system is closed for phosphorus with total concentration *P*_*tot*_.
2. The P:C ratios in the producers vary, while the grazer maintains a constant P:C ratio, *θ*_*y*_. Producer growth depends on minimum requirements *q*_*x*_ and *q*_*w*_, and uptake decreases as the quotas approach 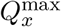 and 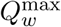.
3. All phosphorus in the system is divided into five pools: phosphorus in the grazer, phosphorus in the two producers, and dissolved phosphorus in the benthic and pelagic zones.

The quota bounds describe the intended biological range. Their preservation by the differential equations requires separate consideration, especially when light varies with time.

Under these assumptions, we can write the total phosphorus as

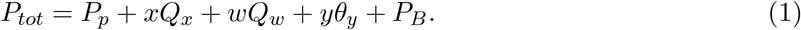

Here, *xQ*_*x*_, *wQ*_*w*_, and *yθ*_*y*_ represent the phosphorus carried by phytoplankton, periphyton, and *Daphnia*, respectively. Table 1 lists the state variables and units.

**Table 1.** State variables and units. The pelagic phosphate pool is obtained from the total phosphorus balance.

| Variable | Description | Units |
| --- | --- | --- |
| $x$ | Phytoplankton biomass | mg C/L |
| $w$ | Periphyton biomass | mg C/L |
| $y$ | *Daphnia* biomass | mg C/L |
| $Q_x, Q_w$ | Producer P:C ratios | mg P/mg C |
| $P_B, P_p$ | Benthic and pelagic phosphate | mg P/L |

### 2.2 Phosphate uptake, growth, and grazing

We first describe phosphate uptake by the producers. The uptake rate is a Michaelis–Menten function of dissolved phosphate, modified by the producer’s P:C ratio:

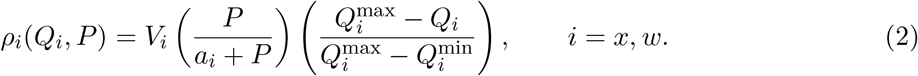

The parameter *V*_*i*_ sets the quota uptake rate, and *a*_*i*_ is the phosphate half-saturation concentration. Uptake increases with available phosphate and decreases as the producer quota approaches its upper value. The lower value 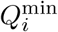 used in uptake normalization is kept distinct from the growth requirement *q*_*i*_ because they play different roles in the formulation.

Next, we consider the rate of change of periphyton and phytoplankton biomass. In words, each producer gains biomass through growth and loses biomass through consumer ingestion. For positive biomass and quota, its specific growth rate is

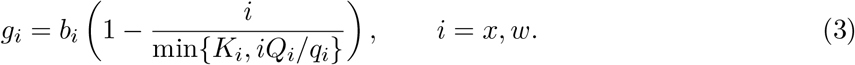

The carrying capacity is determined by whichever resource is limiting: light, represented by *K*_*i*_, or phosphorus, represented by *iQ*_*i*_*/q*_*i*_. For *i >* 0, it is equivalent to *g*_*i*_ = *b*_*i*_[1 − max{*i/K*_*i*_, *q*_*i*_*/Q*_*i*_*}*].

The consumer feeds on both producers with a common ingestion denominator, *c* + *x* + *w*. The P:C ratio of the combined food is (*xQ*_*x*_ + *wQ*_*w*_)*/*(*x* + *w*). Thus, for *x* + *w >* 0, the conversion factor for consumer production is

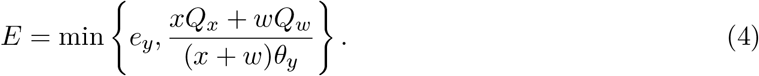

The consumer’s growth depends on both the amount of food and its phosphorus content. The parameters *α, c, e*_*y*_, and *d*_*y*_ denote the maximal ingestion rate, ingestion half-saturation biomass, maximal production efficiency, and consumer loss rate, respectively.

### 2.3 Quota view

Putting these processes together, we obtain six differential equations:

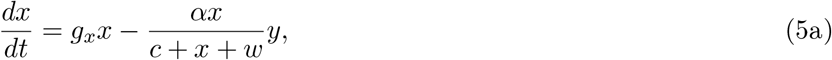

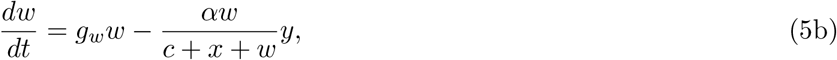

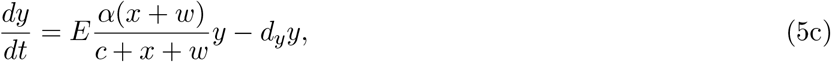

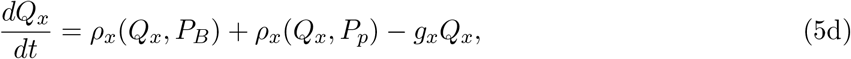

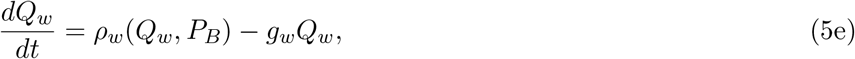

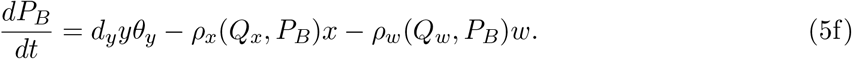

Phytoplankton takes up phosphate from both pools, while periphyton takes up phosphate only from the benthic pool. The quota equations include uptake and growth dilution. Consumer death returns phosphorus to the benthic pool; dietary phosphorus not retained in consumer production returns to the pelagic pool, as shown below. These nutrient pathways are model assumptions. Producer mortality, immigration, and external nutrient loading are not included as separate processes.

Observe that we can reduce the system to six differential equations by assuming that total phosphorus is constant and solving for *P*_*p*_. From Eq. (1),

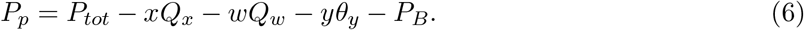

This expression tracks the phosphate remaining in the pelagic zone. We assume positive 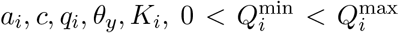, nonnegative *b*_*i*_, *V*_*i*_, *α, d*_*y*_, and 0 ≤ *e*_*y*_ ≤ 1. The same consumer loss rate *d*_*y*_ appears in consumer mortality and benthic phosphorus return.

### 2.4 Nutrient view and phosphorus accounting

Instead of considering the P:C ratios of periphyton and phytoplankton, we can keep track of their phosphorus contents. Let *P*_*x*_ = *xQ*_*x*_ and *P*_*w*_ = *wQ*_*w*_. Applying the product rule to Eq. (5) gives

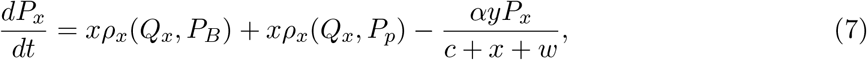

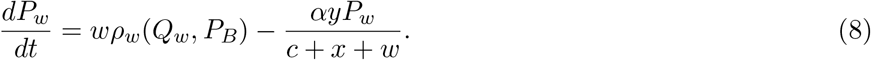

The growth and growth-dilution terms cancel. For positive producer biomasses, *Q*_*x*_ = *P*_*x*_*/x* and *Q*_*w*_ = *P*_*w*_*/w* recover the quota view. Thus, the two formulations are equivalent on this domain. The change of variables is not invertible when a producer biomass is zero. The short calculation is given in Appendix A.

Differentiating Eq. (6) gives the pelagic phosphate balance:

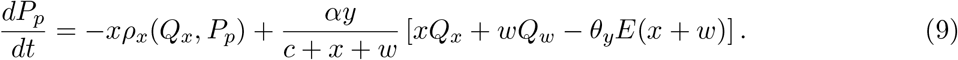

In words, pelagic phosphate decreases through phytoplankton uptake and increases through phosphorus ingested but not retained by the consumer. By Eq. (4), the expression in brackets is nonnegative for nonnegative food biomass and quotas. Summing the producer, consumer, and dissolved phosphorus balances gives *dP*_*tot*_*/dt* = 0. This checks the nutrient accounting of the symbolic model; numerical conservation must also be checked in any implementation.

### 2.5 Light-dependent seasonality

Next, we add light-dependent seasonality to the base model. The base model assumes that periphyton and phytoplankton receive constant light. However, light varies with the time of year. Following the seasonal carrying-capacity approach in [11], we replace *K*_*i*_ by

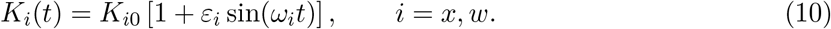

Here, *K*_*i*0_ is the average carrying capacity, *ε*_*i*_ is the amplitude of seasonal forcing, and *ω*_*i*_ is the angular frequency. The magnitude of the perturbation is *ε*_*i*_*K*_*i*0_, and the period is 2*π/ω*_*i*_ for *ω*_*i*_ *>* 0. When *ε*_*i*_ = 0, this light-seasonality model returns the base model. Only the carrying capacities in Eq. (3) change.

For *K*_*i*0_ *>* 0, we require 0 ≤ *ε*_*i*_ *<* 1 to keep the carrying capacities strictly positive. This restriction alone does not guarantee the intended quota bounds. When biomass exceeds the instantaneous carrying capacity, *g*_*i*_ can be negative. At 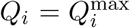, uptake is zero and −*g*_*i*_*Q*_*i*_ can then increase the quota above its upper value. Theorem 3.2 gives a sufficient phosphorus-budget condition that prevents this upper-quota escape and establishes boundedness and positive invariance. The result does not establish persistence.

## 3 Boundedness

With any model, we must be confident that its solutions make sense biologically. Therefore, if we can show that our solutions always remain in a biologically relevant region, then we can be certain that the model respects these biological constraints. Figure 1 illustrates the boundary argument used below. So we prove the following theorem.

**Figure 1.**
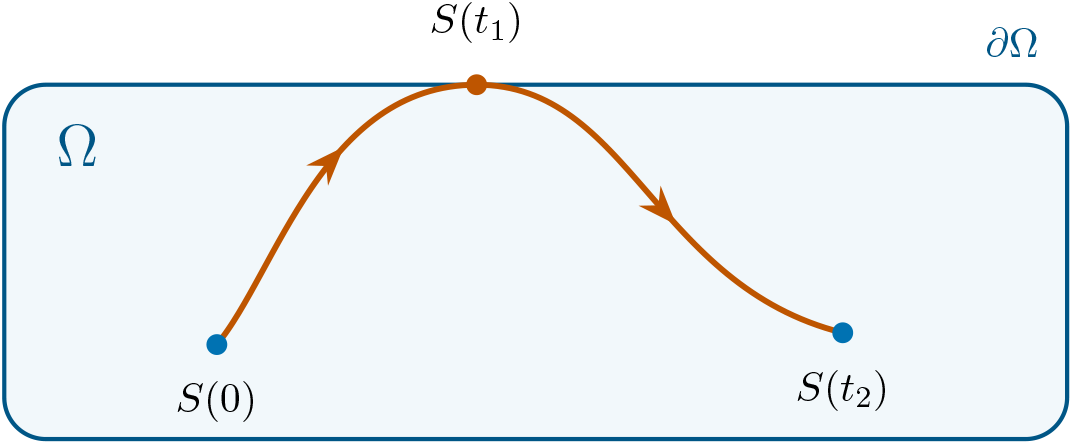
Schematic of the boundary argument. A solution starting in Ω cannot cross outward through ∂Ω when the vector field is inward or tangent at every boundary point. The illustrated contact and return explain the proof idea; they are not a computed trajectory or a claim that every solution reaches the boundary.

At zero producer biomass we use the continuous extension *g*_*i*_ = *b*_*i*_[1 − max {*i/K*_*i*_, *q*_*i*_*/Q*_*i*_*}*]. We also write the full consumer production term as

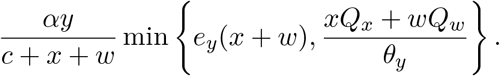

These expressions agree with the quota model for positive producer biomasses and remain defined when either or both producers are absent. Throughout this section, we assume 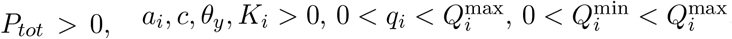, nonnegative *b*_*i*_, *V*_*i*_, *α, d*_*y*_, and 0 ≤ *e*_*y*_ ≤ 1.

### Theorem 3.1

(Boundedness of the base system). *Solutions to the base system with initial conditions in the set*

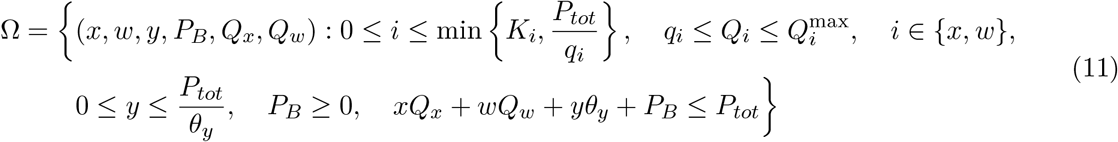

*will remain in* Ω *for all t ≥* 0. *Each such initial condition gives a unique solution defined for all t ≥* 0. *Biomasses that are initially positive remain positive at every finite time. If y*(0) *< P*_*tot*_*/θ*_*y*_, *then y*(*t*) *< P*_*tot*_*/θ*_*y*_ *at every finite time as well*.

The idea of the proof is to show that if a solution *S*(*t*) starts inside our region Ω and reaches its boundary, then it must either stay on the boundary or turn back into the interior of Ω. Figure 1 illustrates this boundary argument.

*Proof*. Let *S*(*t*) = (*x*(*t*), *w*(*t*), *y*(*t*), *P*_*B*_(*t*), *Q*_*x*_(*t*), *Q*_*w*_(*t*)) be a solution to the quota model. We prove the theorem by proving that the solution cannot cross each individual boundary.

To make this boundary argument precise, first regard *P*_*p*_ as a seventh state with differential equation (9). Write

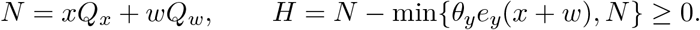

The inequality holds whenever producer biomasses and quotas are nonnegative. In this notation the extended pelagic equation is

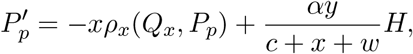

which remains defined when both producers are absent. The extended right-hand side is locally Lipschitz on an open neighborhood of the region considered below: the quotas and uptake denominators are separated from zero, and minima and maxima of locally Lipschitz functions remain locally Lipschitz. Hence solutions exist locally and are unique, including at limitation switches. We first check the rectangular region

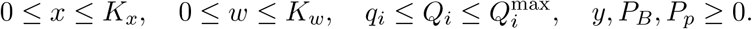

At every boundary face the corresponding derivative points inward or is zero. The following cases therefore establish positive invariance of this region on the maximal interval of existence, including initial conditions on its boundary.

### Case 1

*Q*_*x*_ = *q*_*x*_. Since *x* ≤ *K*_*x*_, we have *g*_*x*_ = *b*_*x*_[1 − max{*x/K*_*x*_, 1}] = 0. Therefore,

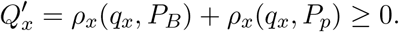

Note that the uptake functions are nonnegative in this region. So the solution cannot cross the boundary *Q*_*x*_ = *q*_*x*_.

### Case 2

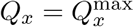. Both phytoplankton uptake terms vanish. Note that *x* ≤ *K*_*x*_ and 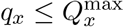, so *g*_*x*_ ≥ 0. Therefore,

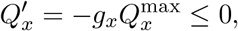

and the solution cannot cross the boundary 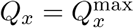.

### Case 3

*Q*_*w*_ = *q*_*w*_. Similarly, *g*_*w*_ = 0, and the periphyton equation gives

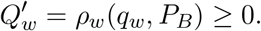

Periphyton takes up phosphate only from the benthic pool. So the solution cannot cross the boundary *Q*_*w*_ = *q*_*w*_.

### Case 4

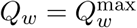. The benthic uptake term vanishes and *g*_*w*_ ≥ 0. Therefore,

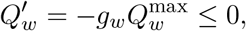

and the solution cannot cross the boundary 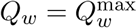.

### Case 5

*y* = 0. The consumer equation gives *y* = 0. Thus an initially absent consumer remains absent, and the solution cannot cross into negative consumer biomass.

### Case 6

*x* = 0. The extended producer equation gives *x* = 0. Thus an initially absent phytoplankton population remains absent, and the solution cannot cross into negative phytoplankton biomass.

### Case 7

*x* = *K*_*x*_. Since *Q*_*x*_ ≥ *q*_*x*_, we have *g*_*x*_ = 0 and

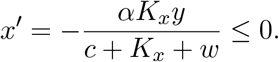

Therefore, the solution cannot cross the boundary *x* = *K*_*x*_. The additional phosphorus bound *x* ≤ *P*_*tot*_*/q*_*x*_ will follow from the total phosphorus balance below.

### Case 8

*w* = *K*_*w*_. Similarly, *g*_*w*_ = 0 and

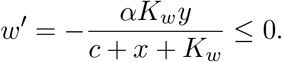

Therefore, the solution cannot cross the boundary *w* = *K*_*w*_.

### Case 9

*w* = 0. The extended producer equation gives *w* = 0. Thus an initially absent periphyton population remains absent, and the solution cannot cross into negative periphyton biomass.

### Case 10

*P*_*p*_ = 0. This is the boundary *xQ*_*x*_ + *wQ*_*w*_ + *yθ*_*y*_ + *P*_*B*_ = *P*_*tot*_ after imposing the phosphorus balance. Pelagic uptake vanishes here, and Eq. (9) gives

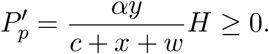

Equivalently, the derivative of *xQ*_*x*_ + *wQ*_*w*_ + *yθ*_*y*_ + *P*_*B*_ on this boundary is −*αyH/*(*c* + *x* + *w*) ≤ 0. Consumer mortality and benthic phosphorus return cancel exactly; the remaining term is the dietary phosphorus not retained by the consumer. Therefore, the solution cannot cross this boundary.

### Case 11

*P*_*B*_ = 0. Both benthic uptake terms vanish, so

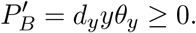

Therefore, the solution cannot cross the boundary *P*_*B*_ = 0.

Summing the phosphorus balances now gives

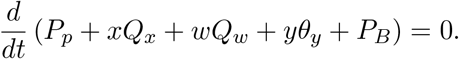

Choose *P*_*p*_(0) = *P*_*tot*_ − *x*(0)*Q*_*x*_(0) − *w*(0)*Q*_*w*_(0) − *y*(0)*θ*_*y*_ − *P*_*B*_(0) ≥ 0. The conserved total is then *P*_*tot*_, and the seven-state solution is equivalent to the six-state system. Since the phosphorus in either producer cannot exceed the total phosphorus in the system, we have *xQ*_*x*_ ≤ *P*_*tot*_ and *wQ*_*w*_ ≤ *P*_*tot*_.

Together with *Q*_*i*_ ≥ *q*_*i*_, these imply *i* ≤ *P*_*tot*_*/q*_*i*_. Similarly, *y* ≤ *P*_*tot*_*/θ*_*y*_ and *P*_*B*_, *P*_*p*_ ≤ *P*_*tot*_. Thus the solution remains in Ω.

The region Ω is compact and remains separated from every zero denominator. The local solution can therefore be continued for all *t* ≥ 0.

Finally, throughout Ω we have *g*_*i*_ ≥ 0 and

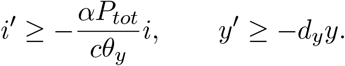

By the standard comparison argument,

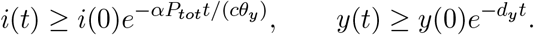

These bounds are strictly positive when the corresponding initial biomass is positive. To retain the strict consumer upper bound, let *R* = *P*_*tot*_ − *θ*_*y*_*y*. Since *N* ≤ *R* and consumer production is limited by dietary phosphorus,

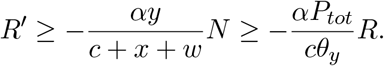

Hence 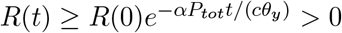 if *R*(0) *>* 0, which proves the last assertion.

### 3.1 Boundedness with light-dependent seasonality

For the seasonal model, the upper-quota argument requires a further condition. Define

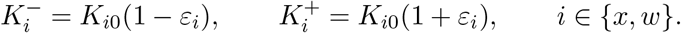

The phosphorus budget supplies the needed bound when

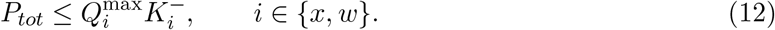

This condition is sufficient; we do not assert that it is necessary.

#### Theorem 3.2

(Boundedness of the light-dependent seasonality model). *Assume the parameter conditions of Theorem 3*.*1, with K*_*i*_ *replaced by Eq*. (10), *K*_*i*0_ *>* 0, 0 ≤ *ε*_*i*_ *<* 1, *and Eq*. (12). *Solutions to the light-dependent seasonality model with initial conditions in the set*

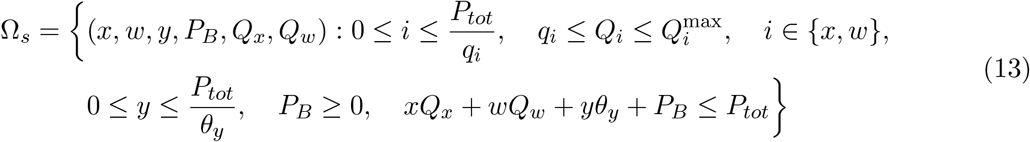

*will remain in* Ω_*s*_ *for all t* ≥ 0. *Each such initial condition gives a unique solution defined for all t* ≥ 0. *Initially positive biomasses remain positive at finite times, and the strict consumer bound is retained if it holds initially. Moreover*,

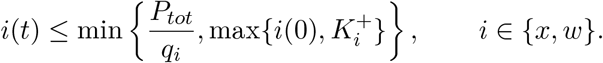

*Proof*. As in the proof of Theorem 3.1, let *S*(*t*) = (*x*(*t*), *w*(*t*), *y*(*t*), *P*_*B*_(*t*), *Q*_*x*_(*t*), *Q*_*w*_(*t*)) be a solution to the quota model. We prove the theorem by proving that the solution cannot cross each individual boundary.

The bounds *i* ≤ *P*_*tot*_*/q*_*i*_ and *y* ≤ *P*_*tot*_*/θ*_*y*_ follow from nonnegative biomasses and pools, the lower-quota bounds, and the phosphorus budget. Thus it suffices to check the nonnegative biomass and benthic-pool faces, both quota faces, and the budget face.

At *Q*_*i*_ = *q*_*i*_, the seasonal growth rate satisfies

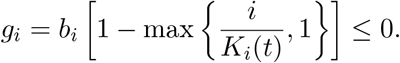

Uptake is nonnegative, so 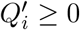. At 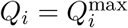, the phosphorus budget and Eq. (12) imply

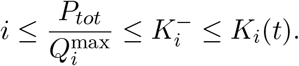

Both arguments of the maximum in *g*_*i*_ are therefore at most one, giving *g*_*i*_ ≥ 0. Uptake vanishes at this quota, and hence 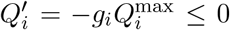. This supplies the upper-quota boundary argument for the seasonal model.

At zero biomass, the corresponding biomass derivative is zero. At *P*_*B*_ = 0, 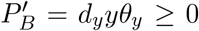. At the budget face *P*_*p*_ = 0, the same phosphorus-accounting calculation as in Case 10 gives 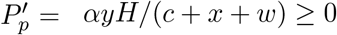. None of these signs depends on the carrying capacities being constant.

For completeness, these face conditions also apply where several boundaries meet. The independent constraints are nonnegative biomasses and *P*_*B*_, the quota intervals, and the budget inequality. At a boundary point one can move every active quota into its interval and every zero biomass or pool toward positive values. If the budget face is also active, its total equals *P*_*tot*_ *>* 0, so at least one of *x, w, y, P*_*B*_ is positive. Decreasing the positive biomass and pool coordinates sufficiently makes the budget decrease while retaining the other inward directions. Thus there is a direction strictly inward to every active constraint, and the tangent cone is given by their inward half-spaces. The locally Lipschitz vector field satisfies these inequalities at every time. The boundary conditions therefore establish positive invariance, including at corners. Compactness and the positive lower bounds on *K*_*i*_(*t*) and the denominators give global existence and uniqueness.

The biomass bound follows by the standard comparison argument, since

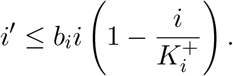

For positivity, set

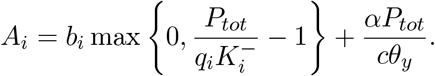

Throughout Ω_*s*_, *i* ≥ − *A*_*i*_*i* and *y* ≥ −*d*_*y*_*y*. Therefore 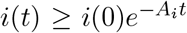 and 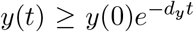. The same argument for *R* = *P*_*tot*_ − *θ*_*y*_*y* used in the base-system proof preserves the strict consumer upper bound.

Condition (12) restricts this result to parameter sets for which the phosphorus budget prevents upper-quota escape. Positive carrying capacities alone do not establish that conclusion. No modification of the growth or uptake functions is used in either theorem.

## 4 Equilibria and local stability

### 4.1 Grazer-free equilibria

We proceed by finding the equilibria with no consumer. These have the general forms

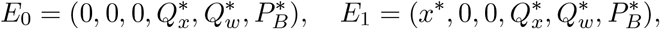

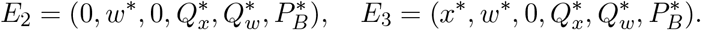

Throughout this calculation, the kinetic parameters are positive and 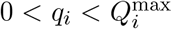, with 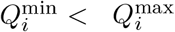. We consider equilibria satisfying *P*_*B*_, *P*_*p*_ ≥ 0 and *q*_*i*_ ≤ *Q*_*i*_ ≤ *Q*_*i*_. For the constant-light model, Theorem 3.1 establishes forward invariance when the complete initial-state conditions in Eq. (11) hold.

The growth expression contains a zero denominator when a producer is absent. For positive quotas we therefore use its equivalent continuous extension

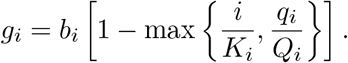

We likewise write the full consumer production term as

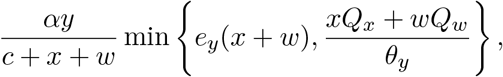

which is defined when both producer biomasses are zero. At *y* = 0 its first-order consumer row is well defined even at the consumer conversion switch; at *x* = *w* = *y* = 0 the production term is of second order and the consumer loss contributes −*d*_*y*_. Producer limitation switches still require separate branch analysis.

An absent producer is on the nutrient-limited branch, since *i/K*_*i*_ = 0 *< q*_*i*_*/Q*_*i*_. Its quota must still satisfy its own differential equation. Consequently, the quotas of absent producers cannot be assigned arbitrary values.

For compactness, let *P* = *P*_*tot*_ and 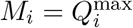. Define

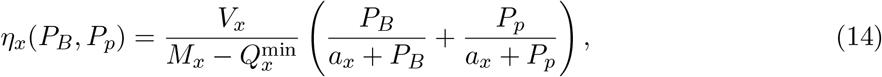

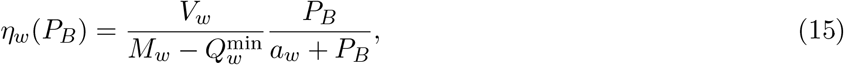

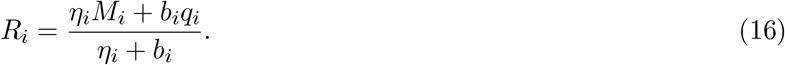

For an absent producer, its quota equation reduces to *η*_*i*_(*M*_*i*_ − *Q*_*i*_) −*b*_*i*_(*Q*_*i*_ − *q*_*i*_) = 0, giving *Q*_*i*_ = *R*_*i*_. In particular, *R*_*i*_ = *q*_*i*_ when its accessible dissolved phosphate is zero, and *q*_*i*_ *< R*_*i*_ *< M*_*i*_ when accessible phosphate is positive.

For a present producer at a grazer-free equilibrium, its biomass equation gives *g*_*i*_ = 0. We write its feasible biomass–quota pairs as

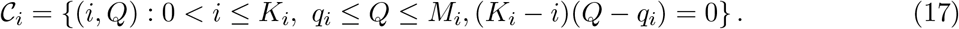

Thus a resident has either its light-limited biomass *i* = *K*_*i*_, its growth requirement *Q* = *q*_*i*_, or both at the switching point. Its quota equation also requires *η*_*i*_(*M*_*i*_ − *Q*_*i*_) = 0.

These conditions give the following complete families of grazer-free equilibria.

1. With both producers absent,

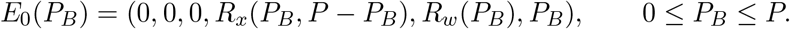 Thus even the organism-free equilibrium is a family parameterized by the split of dissolved phosphate between habitats.
2. With phytoplankton alone and *P*_*B*_ + *P*_*p*_ *>* 0,

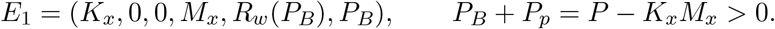 If *P*_*B*_ = *P*_*p*_ = 0, then

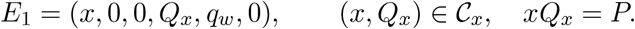 The latter gives *x* = *P/q*_*x*_, *Q*_*x*_ = *q*_*x*_ when 0 *< P* ≤ *K*_*x*_*q*_*x*_, or *x* = *K*_*x*_, *Q*_*x*_ = *P/K*_*x*_ when *K*_*x*_*q*_*x*_ ≤ *P* ≤ *K*_*x*_*M*_*x*_.
3. With periphyton alone and *P*_*B*_ *>* 0,

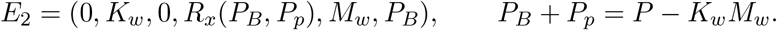 If *P*_*B*_ = 0, then

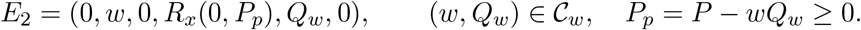 Unlike phytoplankton, periphyton cannot take up pelagic phosphate. A periphyton-only equilibrium can therefore retain *P*_*p*_ *>* 0 without forcing its quota to the maximum.
4. With both producers present and *P*_*B*_ *>* 0,

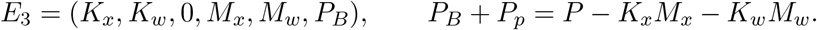 If *P*_*B*_ = 0 and *P*_*p*_ *>* 0, then

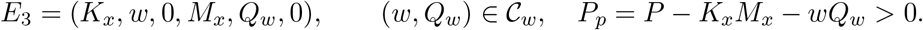 Finally, if *P*_*B*_ = *P*_*p*_ = 0, then

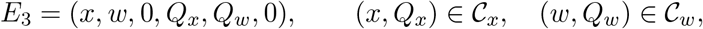

with *xQ*_*x*_ + *wQ*_*w*_ = *P*. This last family includes the nutrient/nutrient, nutrient/light, light/nutrient and light/light cases.

All displayed habitat phosphate concentrations must be nonnegative. The conditions follow directly from the zero biomass and quota derivatives. They also make each benthic uptake term vanish, so the benthic phosphate derivative is zero. Substitution verifies all six equations.

The two consumer conversion regimes give the same grazer-free equilibrium locations because consumer production is multiplied by *y* = 0. They need not give the same consumer invasion rate. Likewise, these families should not be treated as a finite set of isolated points: a smooth family supplies a neutral direction, and switching points require a separate analysis. Consumer-present equilibria with one or two producers are not classified by this grazer-free catalogue.

### 4.2 Local stability of the grazer-free equilibria

We now investigate the stability of the boundary equilibria using the Jacobian of the six governing equations. All spectra below count eigenvalues with multiplicity and apply within a smooth producer-limitation regime. At a switching boundary, the adjacent one-sided Jacobians must be considered separately.

Let *η*_*x*_ and *η*_*w*_ denote the nonnegative uptake coefficients used in the equilibrium conditions, and let *R*_*i*_ denote the corresponding quota of an absent producer. We write

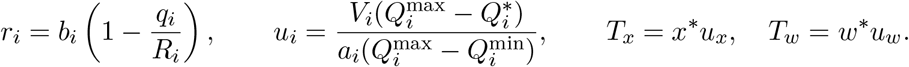

Here *u*_*i*_ is the derivative of uptake with respect to dissolved phosphate at zero phosphate. In the feasible quota range, *u*_*i*_ ≥ 0. At a grazer-free equilibrium with *S*^∗^ = *x*^∗^ + *w*^∗^ *>* 0, the consumer invasion eigenvalue is

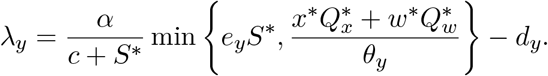

At *E*_0_, the continuous extension of the consumer growth product gives *λ*_*y*_ = −*d*_*y*_.

The consumer row of the Jacobian at *y*^∗^ = 0 has no off-diagonal entries. Thus, if *J*_5_ denotes the block for (*x, w, Q*_*x*_, *Q*_*w*_, *P*_*B*_), the characteristic polynomial factors as

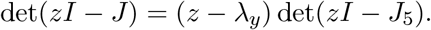

The resulting six eigenvalues are listed in Table 2. The factors were checked by exact symbolic computation. In particular, when *P*_*B*_ = *P*_*p*_ = 0, the benthic and pelagic uptake derivatives cancel in 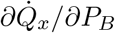. The benthic phosphate eigenvalue is then − (*T*_*x*_ + *T*_*w*_). For either smooth limitation regime of a present phytoplankton population, its biomass–quota block contributes −*b*_*x*_ and −*T*_*x*_; the corresponding periphyton block contributes −*b*_*w*_ and zero. These observations give the same factorization for all four producer regimes of *E*_3_ at zero dissolved phosphate. The absent-producer quota derivatives give the *E*_1_ and *E*_2_ specializations in the table.

**Table 2.** Spectra of feasible grazer-free equilibrium families. The equilibrium conditions and quota constraints must also hold. Each row lists six eigenvalues, including repeated roots.

| Family and dissolved phosphate | Eigenvalues |
| --- | --- |
| $E_0$ | $r_x, r_w, -d_y, -(\eta_x + b_x), -(\eta_w + b_w), 0$ |
| $E_1, P_B + P_p > 0$ | $-b_x, r_w, \lambda_y, -\eta_x, -(\eta_w + b_w), 0$ |
| $E_1, P_B = P_p = 0$ | $-b_x, -b_w, \lambda_y, -T_x, -T_x, 0$ |
| $E_2, P_B > 0$ | $r_x, -b_w, \lambda_y, -(\eta_x + b_x), -\eta_w, 0$ |
| $E_2, P_B = 0, P_p \geq 0$ | $r_x, -b_w, \lambda_y, -(\eta_x + b_x), -T_w, 0$ |
| $E_3, P_B > 0$ | $-b_x, -b_w, \lambda_y, -\eta_x, -\eta_w, 0$ |
| $E_3, P_B = 0, P_p > 0$ | $-b_x, -b_w, \lambda_y, -\eta_x, -T_w, 0$ |
| $E_3, P_B = P_p = 0$ | $-b_x, -b_w, \lambda_y, -T_x, -(T_x + T_w), 0$ |

#### Theorem 4.1

*For positive total phosphate, the feasible organism-free equilibrium family E*_0_ *is unstable to phytoplankton introduction*.

*Proof*. Since *P*_*B*_ +*P*_*p*_ = *P*_*tot*_ *>* 0, phytoplankton has a positive uptake coefficient *η*_*x*_. Its equilibrium quota is 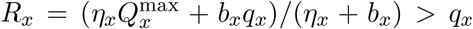. Therefore *r*_*x*_ = *b*_*x*_(1 − *q*_*x*_*/R*_*x*_) *>* 0. The zero eigenvalue reflects the equilibrium family and does not remove this invasion direction.

#### Theorem 4.2

*A feasible phytoplankton-only equilibrium E*_1_ *is unstable if P*_*B*_ *>* 0 *or λ*_*y*_ *>* 0. *When P*_*B*_ = 0 *and λ*_*y*_ ≤ 0, *the eigenvalue test alone does not settle nonlinear stability*.

*Proof*. For *P*_*B*_ *>* 0, *η*_*w*_ *>* 0 and *R*_*w*_ *> q*_*w*_, so *r*_*w*_ *>* 0. A positive *λ*_*y*_ independently gives consumer invasion. At *P*_*B*_ = 0, the absent periphyton quota is *q*_*w*_, giving *r*_*w*_ = 0 rather than *b*_*w*_. The remaining roots in Table 2 are nonpositive when *λ*_*y*_ ≤ 0, and at least one root is zero.

#### Theorem 4.3

*A feasible periphyton-only equilibrium E*_2_ *is unstable if P*_*B*_ + *P*_*p*_ *>* 0 *or λ*_*y*_ *>* 0. *When both dissolved phosphate pools vanish and λ*_*y*_ ≤ 0, *the eigenvalue test alone does not settle nonlinear stability*.

*Proof*. If either dissolved pool is positive, then *η*_*x*_ *>* 0, *R*_*x*_ *> q*_*x*_, and *r*_*x*_ *>* 0. If both pools vanish, *R*_*x*_ = *q*_*x*_ and the phytoplankton invasion eigenvalue is zero. The remaining conclusions follow from Table 2.

#### Theorem 4.4

*A feasible producer-coexistence equilibrium E*_3_ *is unstable to consumer introduction if λ*_*y*_ *>* 0. *Its other eigenvalues are nonpositive in the smooth regimes considered here. For λ*_*y*_ ≤ 0, *the zero eigenvalue prevents a conclusion of local asymptotic stability from the linearization alone*.

*Proof*. Positive parameters and feasible quotas imply *b*_*i*_ *>* 0, *η*_*i*_ ≥ 0, and *T*_*i*_ ≥ 0. All roots in the three *E*_3_ rows of Table 2, apart from *λ*_*y*_, are therefore nonpositive. Every row also contains a zero root.

These results distinguish invasion from persistence. A positive invasion eigenvalue identifies a direction in which a small introduction can grow locally; it does not establish long-term coexistence or guarantee convergence to another equilibrium. Similarly, *λ*_*y*_ = 0 is a candidate consumer invasion boundary, not evidence of a Hopf bifurcation. The bifurcation diagrams must be interpreted together with feasibility, branch conditions, and the computed stability of the corresponding solutions.

## 5 Consumer growth and invasion conditions

The bifurcation diagrams describe how population states change as light availability varies. We first identify the food-quantity and food-quality conditions that govern consumer growth. Let *S* = *x*+*w* be the total producer biomass and *N* = *xQ*_*x*_ + *wQ*_*w*_ the phosphorus carried by the two producers. For *y >* 0 and *S >* 0, Eq. (5c) gives

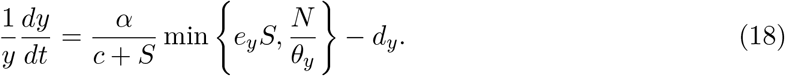

Thus, for *α >* 0, consumer growth is positive precisely when both

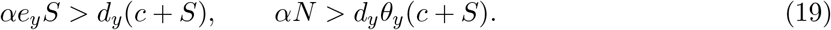

The first inequality is a food-quantity requirement. When *αe*_*y*_ *> d*_*y*_, it can be written as *S > d*_*y*_*c/*(*αe*_*y*_ − *d*_*y*_); if *αe*_*y*_ ≤ *d*_*y*_, positive consumer growth is impossible. The second inequality is the phosphorus requirement for consumer production. Increasing producer biomass can satisfy the first requirement without satisfying the second. This is why light enrichment can support more producer biomass without necessarily supporting greater consumer growth.

At a feasible grazer-free equilibrium with positive producer biomasses, let *S*^∗^ = *x*^∗^ + *w*^∗^ and 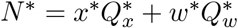. Linearizing the consumer equation in the rare-consumer direction gives

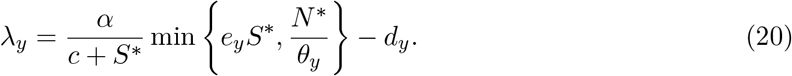

The sign of *λ*_*y*_ determines whether a rare consumer initially increases or decreases at that equilibrium. On a smooth branch of the full system, it is a transverse eigenvalue because the derivatives of the consumer equation with respect to the producer and nutrient variables vanish at *y* = 0. A positive value therefore implies instability to consumer introduction. A negative value describes decay of a rare consumer, but does not establish stability with respect to perturbations of the producers and nutrient pools.

The condition *λ*_*y*_ = 0 identifies a candidate consumer-invasion boundary. It does not identify a Hopf bifurcation: that classification requires an appropriate complex eigenvalue pair and its crossing conditions. The minimum functions also introduce boundaries between active limitation regimes, where a single smooth Jacobian need not apply. We therefore distinguish the equilibrium and periodic branches shown numerically from unverified local bifurcation types. At a coexistence equilibrium, the corresponding consumer balance is

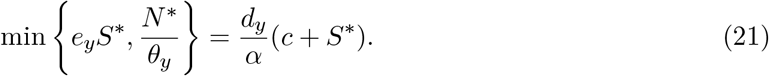

This balance connects the consumer equilibrium branch to the food supplied by both producers.

## 6 Bifurcation analysis and numerical results

### 6.1 Parameters and numerical methods

We vary the common light-dependent carrying capacity, *K*_*x*_ = *K*_*w*_ = *K*, to examine changes in the population dynamics. In the seasonal examples, *K*_0_ denotes the common mean carrying capacity. The parameters used in every figure are listed in Table 3; biological motivation for this parameterization includes [8].

**Table 3.** Parameters used in the recomputed figures. Uptake normalization is specialized to 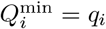; the analytical results retain the more general notation.

| Parameter | Description | Value | Units |
| --- | --- | --- | --- |
| $b_x, b_w$ | Maximal producer growth rates | 1.1, 1.3 | day<sup>-1</sup> |
| $\alpha$ | Maximal consumer ingestion rate | 0.8 | day<sup>-1</sup> |
| $c$ | Ingestion half-saturation biomass | 0.25 | mg C/L |
| $e_y$ | Maximal production efficiency | 0.8 | unitless |
| $d_y$ | Consumer loss rate | 0.25 | day<sup>-1</sup> |
| $\theta_y$ | Consumer constant P:C | 0.03 | mg P/mg C |
| $q_x, q_w$ | Producer minimum growth quotas | 0.0038 | mg P/mg C |
| $Q_x^{\max}, Q_w^{\max}$ | Producer upper quotas | 2.5 | mg P/mg C |
| $Q_x^{\min}, Q_w^{\min}$ | Uptake normalization quotas | 0.0038 | mg P/mg C |
| $a_x, a_w$ | Phosphate half-saturation concentrations | 0.0015 | mg P/L |
| $V_x, V_w$ | Quota uptake rates | 0.2, 0.3 | mg P/(mg C day) |
| $P_{tot}$ | Total phosphorus | 0.03 | mg P/L |

All time series start from (*x, w, y, Q*_*x*_, *Q*_*w*_, *P*_*B*_) = (0.5, 0.5, 0.25, 0.0038, 0.0038, 0.01), with *P*_*p*_(0) = 0.0087 mg P/L. We integrate for 200 days under constant light and 1,000 days under seasonal forcing. For seasonality we use *ω* = 4*π/*365 day^−1^, giving a period of 182.5 days. This is a documented reconstruction choice supported by the available forcing code and trajectory comparisons; the original light-only run configuration was not recovered. These initial states and parameters satisfy the nutrient-budget region and sufficient condition in Theorem 3.2, including its constant-light specialization. This matters for the lower carrying capacities, where initial biomass exceeds *K*_*i*_.

The equations were solved independently using Octave ode45 and Python DOP853 with relative and absolute tolerances 10^−10^. The plotted data use DOP853 with both tolerances tightened to 10^−12^. Across 22 target and diagnostic configurations, the maximum absolute cross-solver difference was 6.21 × 10^−7^ and the maximum change after tolerance refinement was 9.18 × 10^−7^. Independently integrating pelagic phosphate as a seventh state gave maximum total-phosphorus drift 1.45 × 10^−10^. All saved samples are retained without smoothing or clipping; tiny negative dissolved-pool values are at numerical-tolerance scale.

We computed equilibrium and periodic branches with AUTO-07p 0.9.2 using the same parameters and equations. Equilibrium continuation was initialized from refined steady states obtained at *K* = 0.2 and *K* = 0.9. We retained 1,982 feasible equilibrium points, with maximum equation residual 6.48 × 10^−10^. Their stability classifications were checked against the six-state Jacobian spectrum. Periodic continuation used 400 mesh intervals and four collocation points, with tolerance 10^−10^. All 113 retained periodic solutions passed independent one-period integration; maximum closure error was 8.56 × 10^−7^. Six refinements to 800 mesh intervals and four independent Floquet calculations support the displayed periodic family. Infeasible continuation tails and unresolved solver events are excluded.

The displayed branches were computed by continuing solutions of the documented model configuration. They do not exactly reproduce the historical continuation data: the limited recovered numerical points disagree, and the original XPPAUT model and complete continuation settings remain unavailable. The quantitative results below refer to the current computation.

### 6.2 Equilibrium and periodic branches under light enrichment

Consider the bifurcation diagram in Figure 2, where *K* is varied and the three rows show phytoplankton, periphyton, and consumer biomass. Solid and dashed curves distinguish stable and unstable equilibria. The shaded envelopes show the minimum and maximum biomass along the computed stable periodic family.

**Figure 2.**
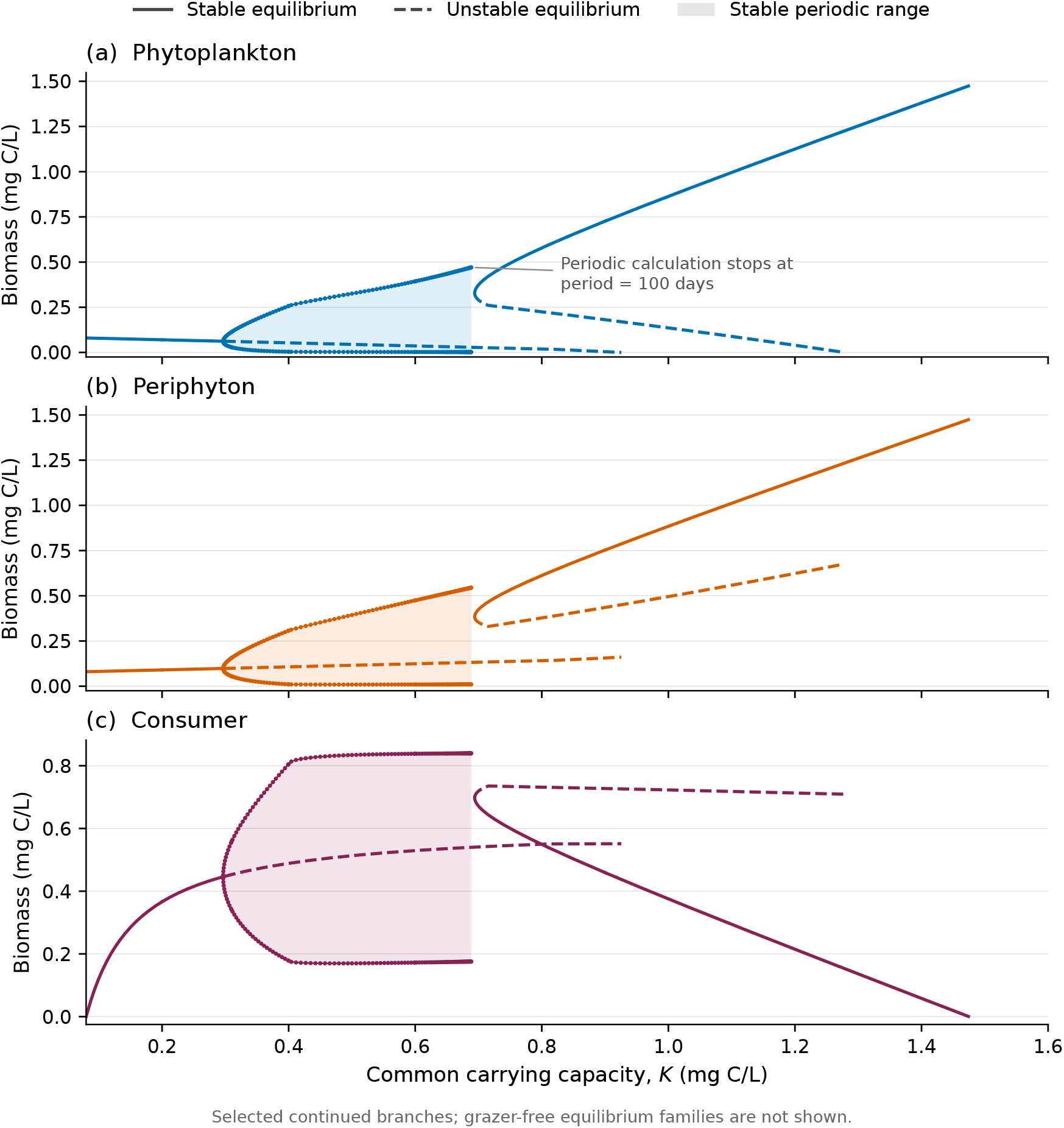
Selected equilibrium and periodic branches computed with AUTO-07p as the common carrying capacity *K* varies. Solid and dashed lines denote stable and unstable equilibria; shaded bands show the minimum and maximum biomass along the stable periodic family. Rows show phytoplankton, periphyton, and consumer biomass. The periodic calculation stops deliberately at period 100 days (*K* ≈ 0.688489); this is not a demonstrated branch termination. Nonisolated grazer-free families are not shown.

The low-biomass equilibrium branch loses stability at a numerical Hopf crossing near *K* = 0.296506. A complex eigenvalue pair crosses the imaginary axis with frequency approximately 0.356434 day^−1^, while the remaining eigenvalues have negative real parts. This crossing occurs away from the limitation switches. The periodic family continued from this point is stable in the retained calculations, and its biomass range broadens as *K* increases. The numerical crossing and periodic calculations do not replace an analytical nondegeneracy proof.

A separate high-biomass equilibrium branch has a numerical fold near *K* = 0.693972, where one real eigenvalue is near zero and the other five have negative real parts. Along its stable segment, producer biomass increases with *K*, while consumer biomass decreases. This response is consistent with the quantity and phosphorus requirements in Eq. (19): greater producer biomass alone is insufficient to maintain consumer production. The descending branch is not a global extinction threshold for all initial conditions.

The periodic calculation stops deliberately when the period reaches 100 days, at *K* ≈ 0.688489. This is a computational cutoff, not a demonstrated termination of the periodic branch. Its proximity to the equilibrium fold does not by itself identify a homoclinic or other global bifurcation. Nonisolated grazer-free equilibrium families are not shown, and the diagram is not an exhaustive search for every branch or attractor.

This branch structure is important for interpreting the response to light. A nearly constant trajectory at one value of *K* does not by itself exclude other solutions. Conversely, the presence of a periodic branch does not imply that all initial conditions approach it. The continuation therefore provides information that a small set of time series cannot supply.

### 6.3 Representative constant-light trajectories

We next examine representative population trajectories under constant light. Figure 3 shows biomass trajectories for *K* = 0.2, 0.4, 0.6, and 0.9. At *K* = 0.2, the trajectories approach nearly constant positive values over the displayed interval. At *K* = 0.4 and *K* = 0.6, the three populations show repeated oscillations. At *K* = 0.9, the trajectories again approach nearly constant positive values. These trajectories sample steady and oscillatory behavior from the stated initial condition and should be interpreted alongside Figure 2.

**Figure 3.**
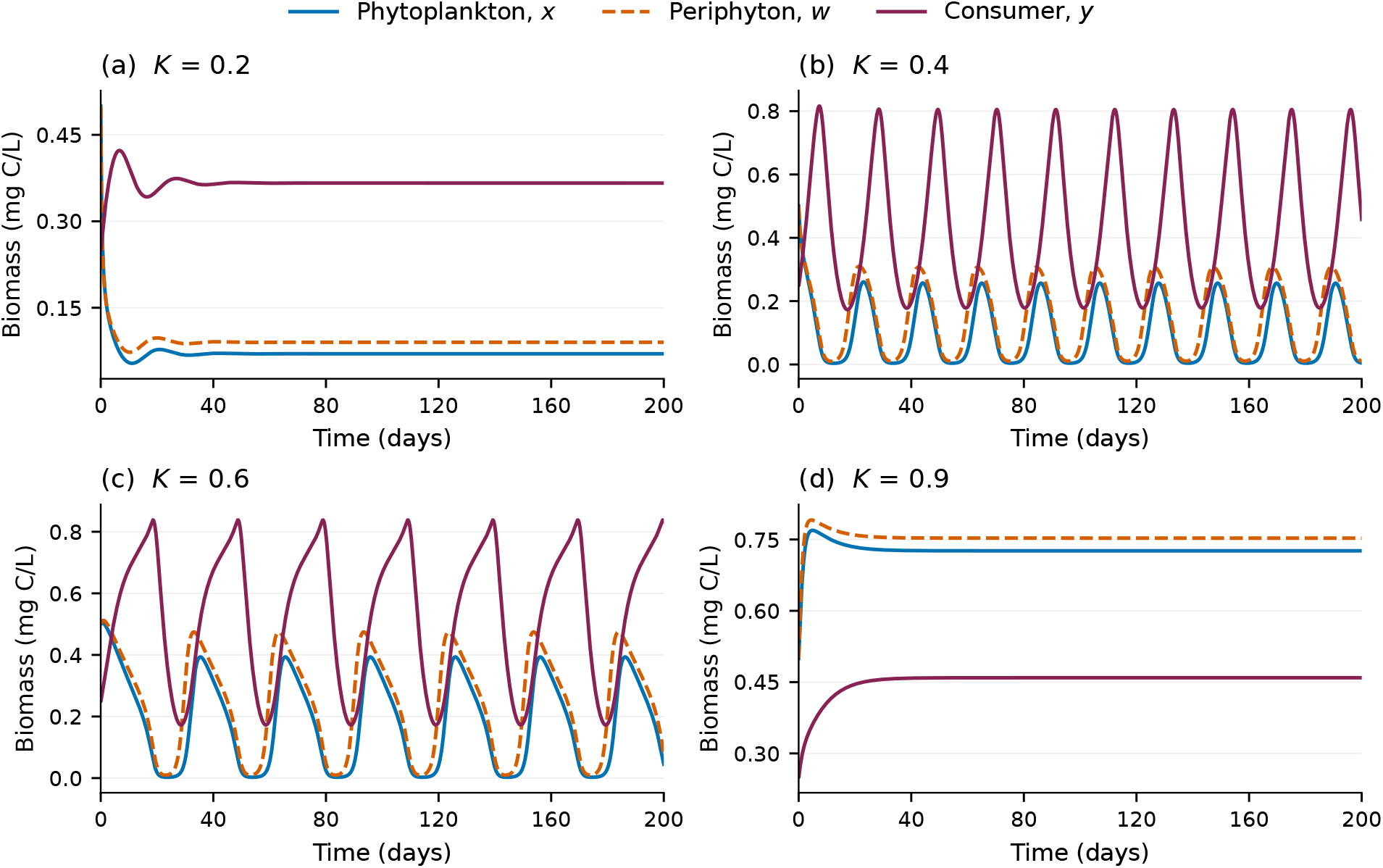
Population trajectories under constant light at *K* = 0.2, 0.4, 0.6, and 0.9 mg C/L. Each panel uses the same initial state and parameters in Table 3. All three populations are shown over the full 200-day interval.

Figure 4 shows the corresponding producer P:C ratios and dissolved phosphate concentrations. These quantities help distinguish the phosphorus content of food from the phosphate remaining in the environment. A higher producer P:C ratio alone does not show that the producer takes up phosphate faster: uptake also depends on dissolved phosphate through Eq. (2), while total uptake depends on producer biomass. Quotas and dissolved phosphate are shown on separate axes with their respective units.

**Figure 4.**
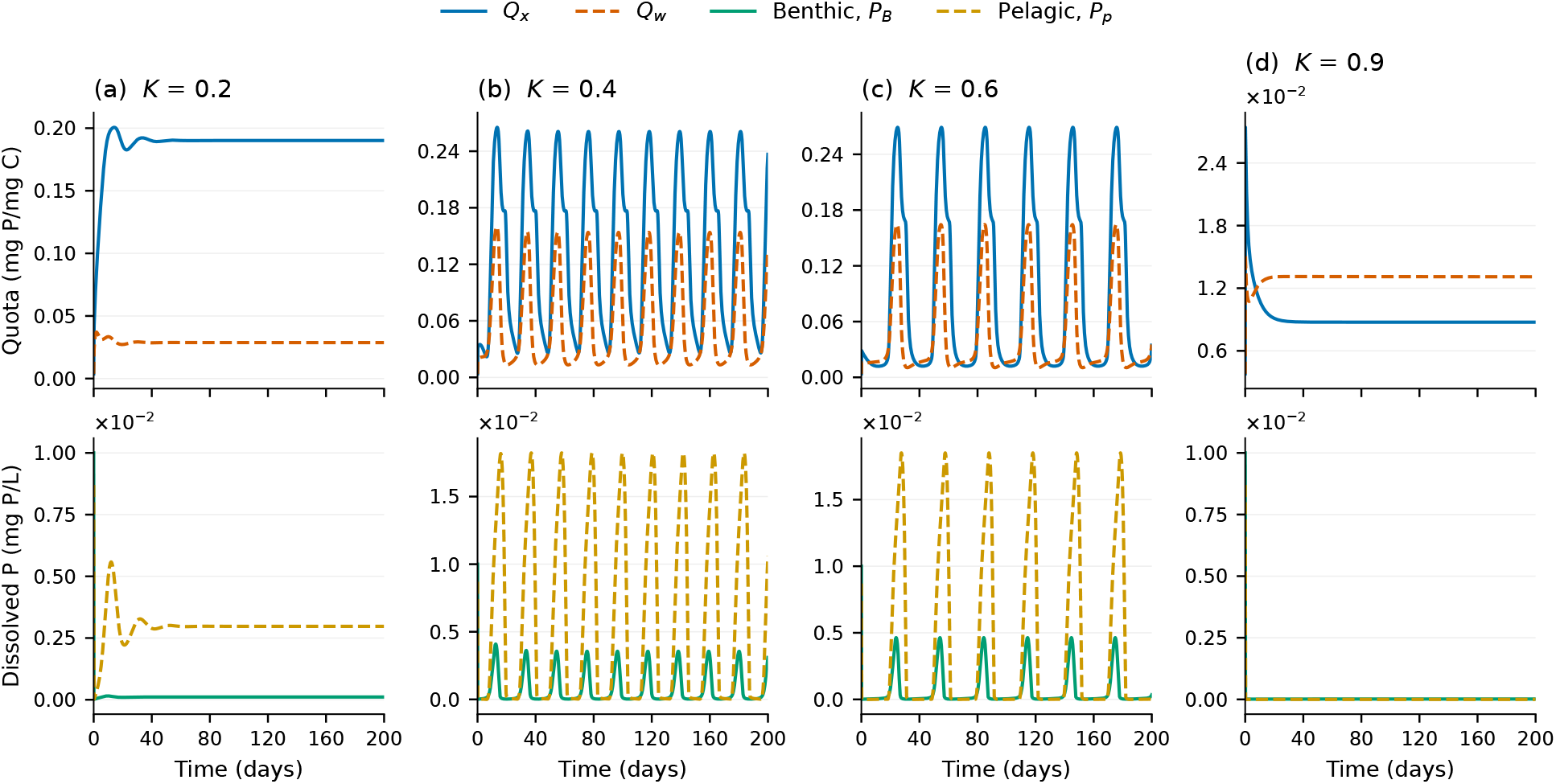
Producer quotas and dissolved phosphate for the constant-light scenarios in Figure 3. Columns correspond to *K* = 0.2, 0.4, 0.6, and 0.9 mg C/L. Quotas (top row) and dissolved phosphate (bottom row) have separate axes and units. Curves are plotted directly from the checked numerical outputs.

### 6.4 Low seasonality

We also vary seasonality from low to high to see its effects on the system’s dynamics. Figure 5 shows the low-seasonality case, *ε* = 0.3, at mean carrying capacities *K*_0_ = 0.25, 0.75, 1, and 2. At *K*_0_ = 2, the *Daphnia* curve declines toward the plotting baseline while producer biomass continues to vary.

**Figure 5.**
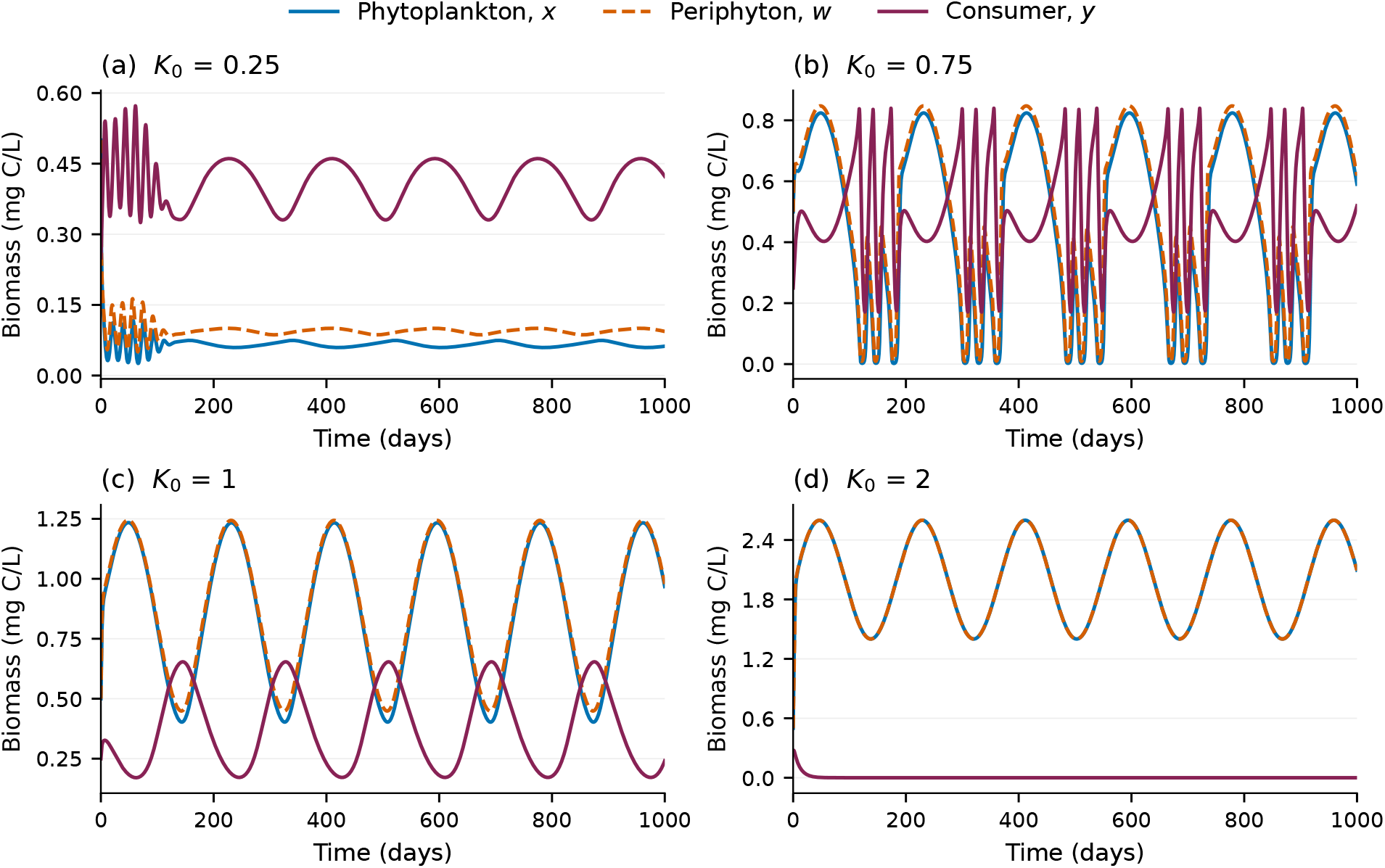
Population trajectories under low seasonality, *ε* = 0.3, at mean carrying capacities *K*_0_ = 0.25, 0.75, 1, and 2 mg C/L. The common forcing period is 182.5 days. The full 1,000-day interval is shown.

Figure 6 shows the producer P:C ratios and dissolved phosphate pools for the same four scenarios. These trajectories provide the food-quality and nutrient-pool context for the biomass changes. The grazer’s P:C ratio is constant in the model.

**Figure 6.**
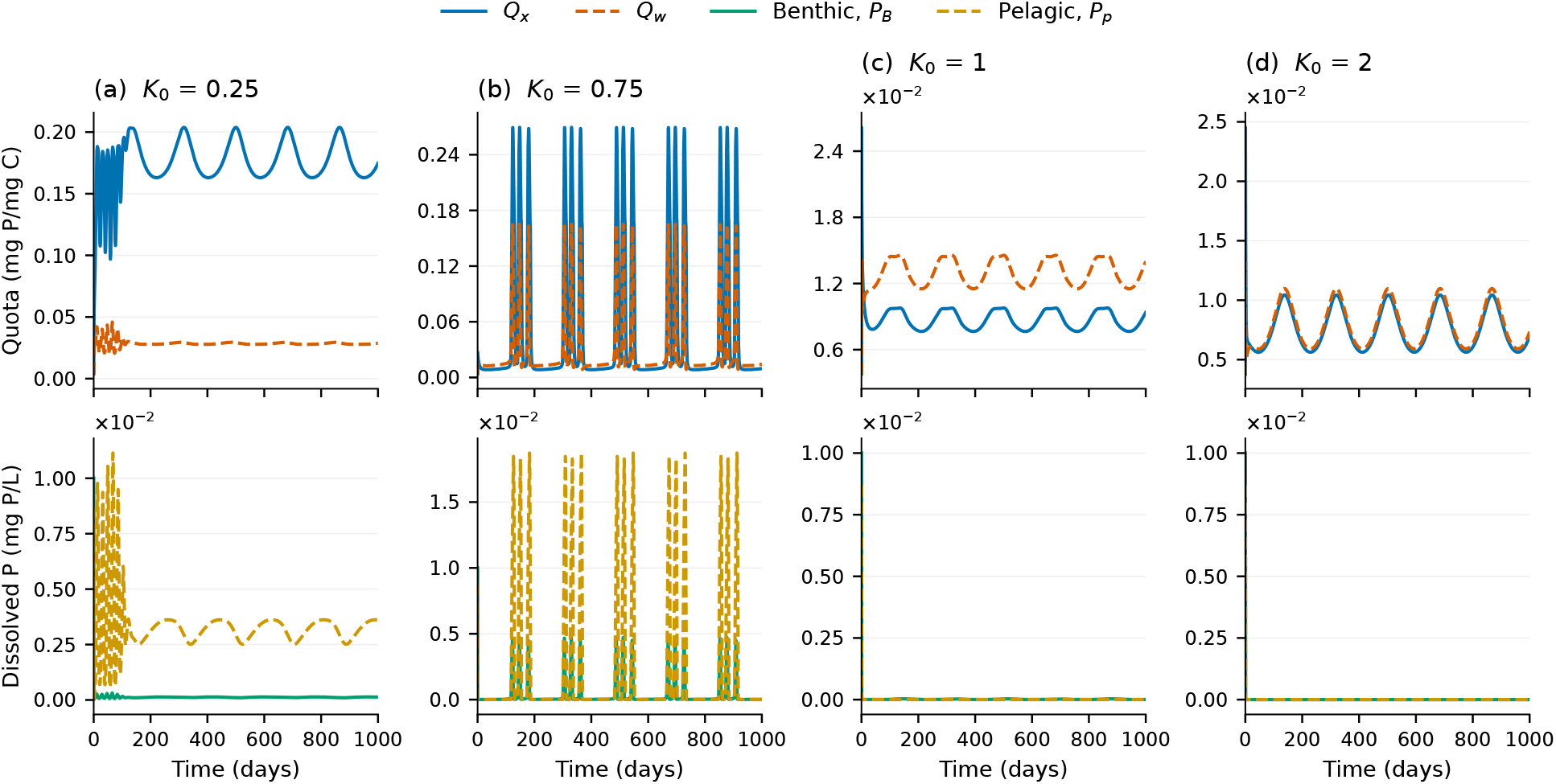
Producer quotas (top row) and dissolved phosphate (bottom row) for the low-seasonality scenarios in Figure 5. Separate axes distinguish food phosphorus content from dissolved-pool concentrations.

### 6.5 High seasonality

At the higher forcing amplitude, *ε* = 0.7, positive consumer peaks occur after the initial decline in the *K*_0_ = 2 scenario (Figure 7). The peaks become smaller over the displayed 1,000 days. Thus, the higher-amplitude example shows brief rebounds; it does not establish survival over repeated years or long-term coexistence.

**Figure 7.**
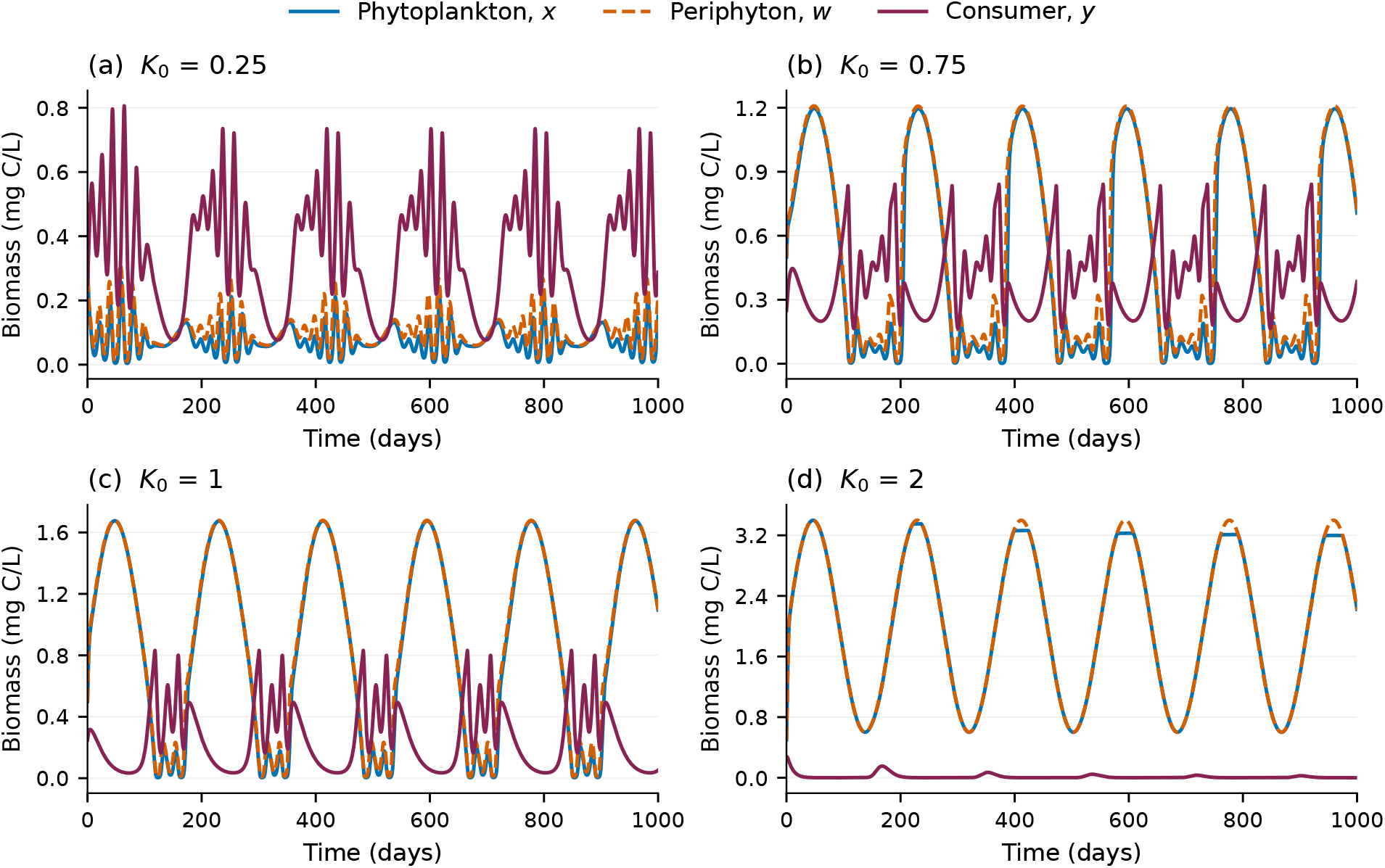
Population trajectories under high seasonality, *ε* = 0.7, with the same mean carrying capacities, forcing period, and initial state as in Figure 5. The consumer rebounds at *K*_0_ = 2 diminish over the displayed interval; these transients do not establish long-term persistence.

The corresponding quota and phosphate trajectories are shown in Figure 8. The light:nutrient hypothesis offers a possible interpretation of the consumer rebounds: periods of lower light may improve producer food quality, allowing *Daphnia* to grow for brief periods. Establishing that explanation requires evaluating the separate quantity and phosphorus growth inequalities along these trajectories. The consumer equation has a factor *y*, so an exactly zero population remains zero. A rebound must therefore arise from a small positive population, rather than recovery from exact extinction.

**Figure 8.**
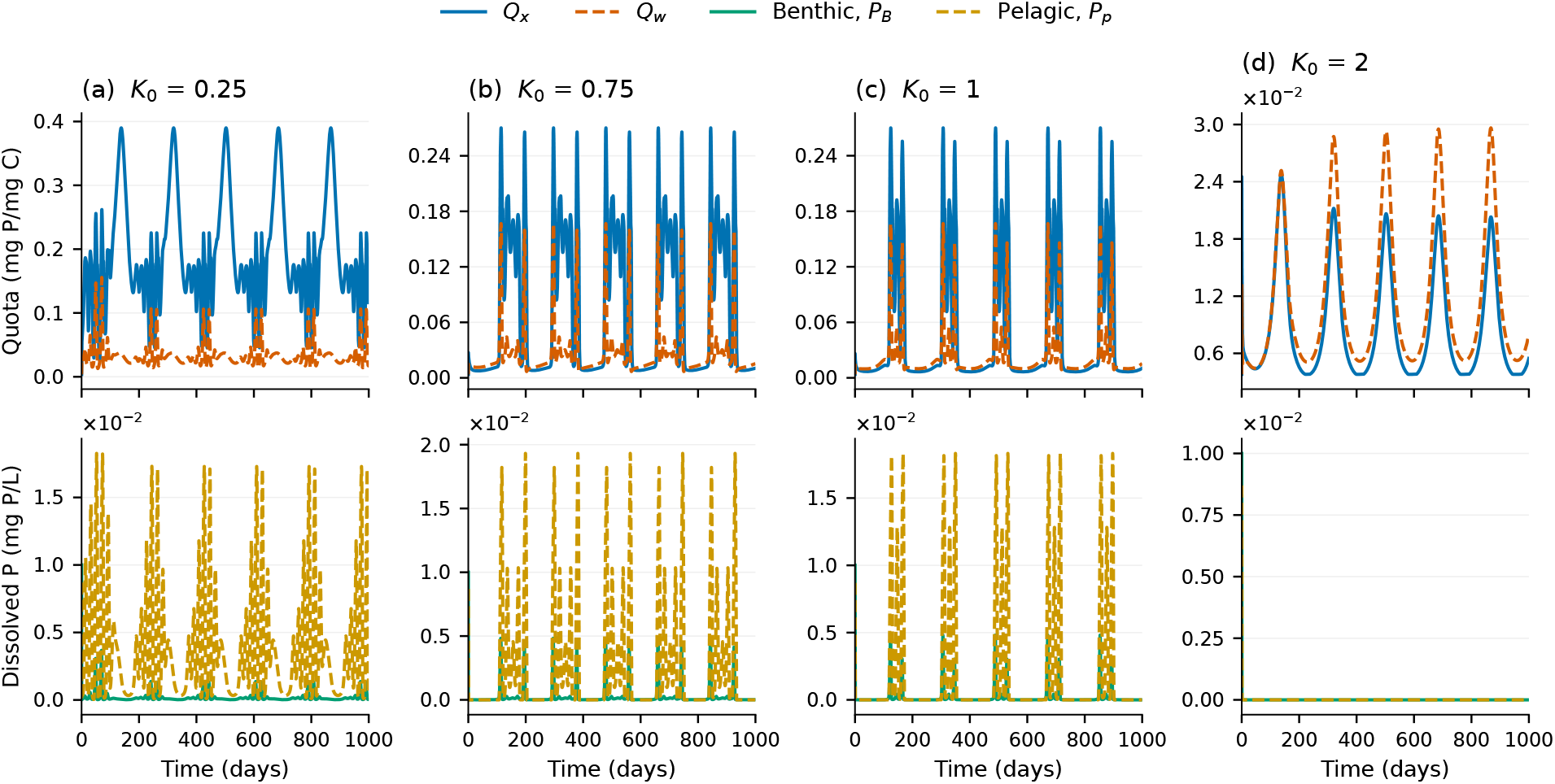
Producer quotas (top row) and dissolved phosphate (bottom row) for the high-seasonality scenarios in Figure 7. All saved samples are retained, with no temporal smoothing or state clipping.

## 7 Discussion

In this study, we constructed a stoichiometric food-web model coupling periphyton and phytoplankton through a shared consumer and separate benthic and pelagic phosphate pools. We then extended the base model by allowing light-dependent carrying capacities to vary with time. The formulation keeps track of both producer biomass and its phosphorus content, allowing us to distinguish the amount of food available from its quality for *Daphnia*.

The bifurcation structure is central to this interpretation. Equilibrium and periodic branches occupy overlapping ranges of light-dependent carrying capacity, and their numerical stability classifications vary along the plotted branches. The selected time series show which behavior is reached in particular examples, while the branch diagrams indicate additional solutions. A few nearly constant or oscillatory trajectories are therefore insufficient to summarize the response to enrichment. A comparable contrast has been observed experimentally: McCauley et al. [33] found large-amplitude *Daphnia*–algal cycles and small-amplitude dynamics in nutrient-rich microcosms from which inedible algae had been excluded. This supports the ecological plausibility of contrasting dynamical regimes. Those microcosms were nutrient enriched, whereas we vary light-dependent carrying capacity. The experiment does not validate our Hopf or fold locations or establish that the same cycles occur generally in lakes.

The descending consumer equilibrium branch illustrates a further ecological distinction. More light can increase the capacity for producer growth without increasing the phosphorus available in food. The consumer growth conditions in Eq. (19) separate these two requirements, and Eq. (20) gives the corresponding local test for consumer introduction at a grazer-free equilibrium. Neither test alone determines the global outcome: the producer–nutrient dynamics and the stability of other branches must also be considered.

This separation of food quantity and quality has support in field experiments. Urabe et al. [34] found that reducing light in a field experiment in a phosphorus-limited lake increased zooplankton herbivore production because algal nutrient content improved relative to carbon. In a field experiment using natural algal communities from six lakes, Striebel et al. [35] found that *Daphnia* growth was highest at intermediate light in oligotrophic and mesotrophic treatments; responses to light were weaker in eutrophic communities. These results are qualitatively consistent with our finding that greater light availability need not improve consumer performance. They concern consumer growth or production, rather than measurements of our equilibrium branches. Hall et al. [5] also found in pond surveys and mesocosms that grazers influence producer stoichiometry, supporting the importance of feedback between consumption and food quality rather than treating quality as a response to light alone.

A more severe food-quality response was observed by Diehl et al. [28]: warming in plankton mesocosms promoted phosphorus-poor algal blooms followed by *Daphnia* collapse despite abundant algal food. This is an experimental analogue of consumer decline in the presence of substantial producer biomass. The forcing was temperature, however, so it supports the underlying stoichiometric mechanism rather than directly validating our light-only model.

Seasonal forcing changes the interpretation again. Instead of approaching a constant environmental state, populations experience repeated changes in producer carrying capacity. Stronger forcing produces brief consumer rebounds in the high-light example, but the peaks diminish over the displayed interval. Periods of positive consumer growth can therefore occur without establishing sustained persistence. A seasonal invasion analysis would require the consumer growth rate along the relevant producer–nutrient trajectory, rather than the constant-equilibrium value used in Eq. (20). Seasonal changes in the limiting resource have also been documented using natural lake food. Müller-Navarra and Lampert [36] combined laboratory growth assays with seasonally collected lake seston and field reproductive observations, finding that *Daphnia* limitation shifted between food quantity and food quality. This supports considering both constraints over a seasonal cycle, but it does not establish phosphorus as the sole cause of food-quality variation or reproduce our diminishing consumer rebounds. The exact rebound sequence and the 182.5-day forcing choice remain model-specific outcomes requiring direct ecological comparison.

The separate phosphate pools make the assumed uptake and recycling pathways explicit. They do not, by themselves, demonstrate a new ecological mechanism. The habitat-coupling studies, shared-grazer models, and seasonal studies discussed above provide different comparisons for this formulation [13, 16, 17, 18, 21]. In particular, the consumer feedback distinguishes it from producer-only competition, while nutrient compartments and feeding assumptions distinguish it from the closest shared-grazer models. A useful next comparison is to retain the same consumer and producer processes while testing a consistent simpler nutrient-pool formulation. Such a comparison would help determine whether the separate pools change consumer growth or the timing of food-quality limitation. Comparisons of stoichiometric model structure and traits in lake ecosystems also provide a useful methodological context [32]. The present figures do not establish nutrient-storage memory of the kind investigated under thermal forcing in [26].

The numerical thresholds in Section 6 apply to the stated parameterization and are not calibrated ecological predictions. The boundedness theorems establish the feasible region under their stated assumptions, but nonlinear dynamics near nonhyperbolic equilibria and the global connections between continued branches require further analysis. In addition, the assumed nutrient access of each producer and the destination of recycled consumer phosphorus need biological justification for the aquatic habitat being represented.

Light is one environmental factor affecting aquatic food webs. Temperature can also affect consumer feeding and producer growth. The temperature-dependent nutritional demands and producer responses identified in recent studies [29, 30, 31] would need to inform such an extension. It would require a consistent parameterization and separate tests of the effects of temperature and food quality. For the current model, the next analytical step is to resolve nonlinear behavior near the nonhyperbolic boundary equilibria, characterize the termination of the periodic branch, and determine when changes in producer food quality lead to sustained consumer growth.

## Acknowledgments

The author thanks Angela Peace for her guidance. This preprint is based on the author’s original dissertation, completed in 2021. The original research, model formulation, calculations, and dissertation text are the author’s work. AI tools assisted with preparing this preprint, including organization, editing, formatting, checking and correcting mathematical derivations, and reproducing numerical figures.

## Data and code availability

The accompanying archive, `preprint-supporting-data-and-code.zip`, contains the simulation configurations, checked time-series outputs, AUTO continuation data, numerical validation reports, and figure-rendering scripts. Symbolic-check scripts reproduce the boundedness identities, Jacobian entries, equilibrium residuals, and local characteristic-polynomial calculations. Software requirements and reproduction commands are documented with these files. The complete original historical continuation configuration remains unavailable.

## A Quota and nutrient-content calculation

For phytoplankton, the product rule gives

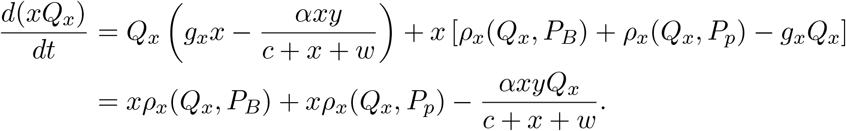

The same calculation for *wQ*_*w*_ gives Eq. (8). Dividing the nutrient contents by positive producer biomasses recovers the quota variables. This establishes the stated change of variables on the positive-biomass domain.

For phosphorus conservation, write *S* = *x* + *w* and *N* = *xQ*_*x*_ + *wQ*_*w*_. On *S >* 0, dietary phosphorus not retained in grazer growth is

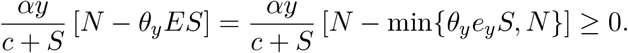

Producer uptake and grazing transfers then cancel in the derivative of Eq. (1). The same *d_y_* in the grazer and benthic equations ensures cancellation of the mortality transfer. These identities establish phosphorus accounting for the stated symbolic system.

## B Full Jacobian matrix

For the state vector *z* = (*x, w, y, Q*_*x*_, *Q*_*w*_, *P*_*B*_)^T^, write the right-hand side of Eq. (5) in the same order as *F* = (*F*_1_, …, *F*_6_)^T^. The full 6 × 6 Jacobian has the form

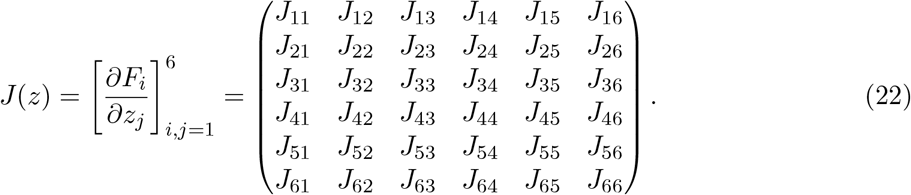

Each row corresponds to one state equation and each column to differentiation with respect to the corresponding component of *z*. The entries, including those that vanish, are given below.

Write *S* = *x* + *w, D* = *c* + *S, N* = *xQ*_*x*_ + *wQ*_*w*_, and *P*_*p*_ = *P*_*tot*_ − *xQ*_*x*_ − *wQ*_*w*_ − *yθ*_*y*_ − *P*_*B*_. Define

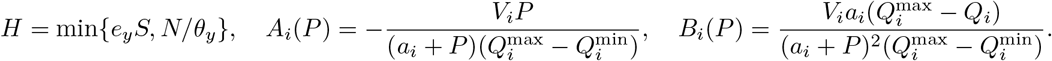

Here *A*_*i*_ and *B*_*i*_ are the uptake partial derivatives with respect to quota and phosphate, respectively. Subscripts *B* and *p* below indicate evaluation at *P*_*B*_ and *P*_*p*_, e.g., *B*_*xp*_ = *B*_*x*_(*P*_*p*_). Also write *r*_*xB*_ = *ρ*_*x*_(*Q*_*x*_, *P*_*B*_) and *r*_*wB*_ = *ρ*_*w*_(*Q*_*w*_, *P*_*B*_). On a strict light-limited producer branch,

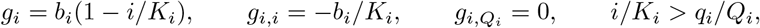

whereas on a strict nutrient-limited branch,

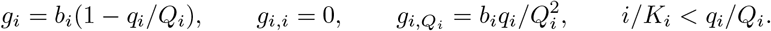

For quantity-limited consumer growth (*e*_*y*_*S < N/θ*_*y*_),

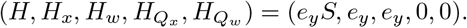

For quality-limited consumer growth (*N/θ*_*y*_ *< e*_*y*_*S*),

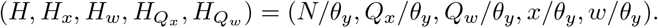

In state order (*x, w, y, Q*_*x*_, *Q*_*w*_, *P*_*B*_), the Jacobian is specified row by row as follows:

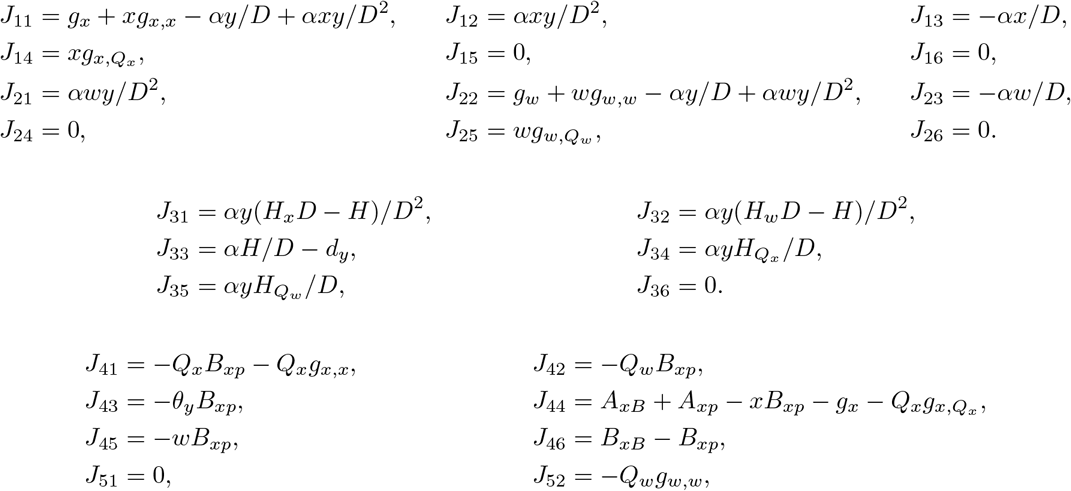

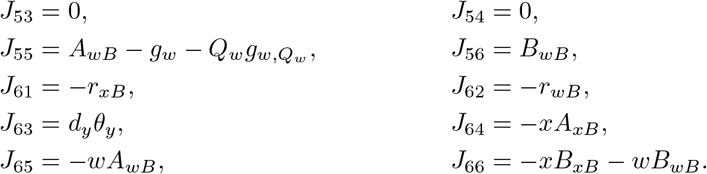

These formulas apply to each of the eight strict combinations of limitation regimes. At a switching surface the neighboring branch Jacobians must be considered separately; selecting one branch at equality does not generally define a classical Jacobian. They assume positive producer biomasses and quotas for the literal growth expressions. Boundary calculations require the explicitly stated extension *g*_*i*_ = *b*_*i*_[1 − max {*i/K*_*i*_, *q*_*i*_*/Q*_*i*_*}*], with *Q*_*i*_ *>* 0, and consumer production *αy* min{*e*_*y*_*S, N/θ*_*y*_*}/D*; this does not make the original quotients defined at zero biomass. A seasonal model has the same state derivatives at fixed time, with *K*_*i*_ = *K*_*i*_(*t*), but constant-equilibrium eigenvalues do not characterize its periodic stability.

